# Key role of auxin cellular accumulation in totipotency and pluripotency acquisition

**DOI:** 10.1101/2022.09.02.505607

**Authors:** Omid Karami, Azadeh Khadem, Arezoo Rahimi, Remko Offringa

## Abstract

Genome editing and *in vitro* based-plant propagation require efficient plant regeneration system. Somatic embryogenesis (SE) or de novo shoot regeneration are two major systems that widely used for plant *in vitro* regeneration. Most SE or shoot regeneration protocols rely on the exogenous application of the synthetic auxin analog 2,4-dichlorophenoxyacetic acid (2,4-D) and naphthylene acetic acid (NAA), whereas the natural auxins indole-3-acetic acid (IAA), 4-chloroindole-3-acetic acid (4-Cl-IAA) or indole-3-butyric acid (IBA) are not or less effective for plant regeneration. Although these synthetic auxins mimics the physiological activity of the main natural auxin IAA in many aspects, there are also clear differences that have been attributed to differences in stability or to different affinities for certain TIR1/AFB-Aux/IAA auxin co-receptor pairs. Here we show that the success of 2,4-D in inducing SE from Arabidopsis is related to ineffectiveness as substrate for auxin efflux, resulting in its intracellular 2,4-D accumulation. Reducing auxin efflux by addition of the auxin transport inhibitor naphthylphthalamic acid (NPA) also allowed natural auxins and other synthetic analogs to induce SE in Arabidopsis with similar efficiencies as 2,4-D. The PIN-FORMED auxin efflux carriers PIN1, PIN2 and the ATP-binding cassette-B auxin transporters ABCB1 and ABCB19 were shown to be partially responsible for the efflux of natural auxins during SE induction. Importantly, all somatic embryos induced in Arabidopsis by IAA in the presence of NPA showed a normal embryo to seedling conversion and subsequent plant development, whereas for the 2,4-D system this was limited to 50-60% of the embryos. We showed that the auxin transport inhibition promotes de novo shoot regeneration capacity from callus induced by 4-Cl-IAA in *Brassica napus*. In addition, we observed a obvious acceleration in shoot bud emerging from callus induced by 4-Cl-IAA than 2,4-D. Based on our data we conclude, that the efficiency of plant propagation can be significantly improved by applying the natural auxins in the presence of the auxin transport inhibitor NPA.

## Introduction

In plant, the plant hormone auxin acts as a master regulator of a wide range of cellular functions. The role of auxin in cellular processes is mainly associated with the level of auxin in plant cells which is determined by de novo auxin biosynthesis, auxin metabolism, auxin homeostasis, and auxin transport (Vanneste and Friml, 2009; Paque and Weijers, 2016).

The auxin is synthesized in all plant cells, but after its biosynthesis in certain cells or tissues, mostly in young developing organs, is transported to sink tissues via the phloem or cell-to-cell transport. A network of auxin influx and efflux plasma membrane-localized proteins transport auxin between cells uniquely in the directional manner is termed polar auxin transport (PAT) (Friml, 2010; Adamowski and Friml, 2015). Three classes of auxin carriers including PIN-FORMED (PIN) proteins (Friml, 2003), ATP-binding cassette-B (ABCB)/P-glycoprotein (PGP) proteins(Geisler *et al*., 2017), AUXIN1/LIKE-AUX1 (AUX/LAX) family members (Péret *et al*., 2012), have been discovered that are responsible for the directionality of auxin flow.

Base on function and subcellular localization, the PIN proteins are categorized into two gropes. The first PIN1-type proteins are auxin efflux carriers that are asymmetrically localized at the plasma membrane and export auxin to the neighboring cell in polar fashion. Expression pattern, dynamics of polar subcellular localization, and abundance of plasma membrane localization of this group of PIN proteins paly important role in direction of auxin export and regulation of auxin gradients (Robert *et al*., 2013; Adamowski and Friml, 2015; Rakusová *et al*., 2016). The second group of PIN proteins that are localized in the endoplasmic reticulum and the nuclear membrane regulate the movement of auxin from the cytoplasm into the lumen of the endoplasmic reticulum and into the nuclear (Mravec *et al*., 2009; Ganguly *et al*., 2010). Second class of auxin transporters are ABCB proteins that facilitate both auxin influx and efflux auxin transport (Geisler *et al*., 2017). ABCB proteins transport auxin against steep auxin gradient, as some ABCB/PGP proteins import auxin into the cytoplasm cells when auxin is low, but they usually export auxin in high auxin level cells ((Yang and Murphy, 2009). Although ABCB are not distributed in at the plasma membrane in a polar fashion (Geisler and Murphy, 2006), by forming complexes with the PINs enhance PAT (Blakeslee *et al*., 2007). The third class of auxin transporters are AUX/LAX proteins that mediate auxin import into the cytoplasm and depends on the cell or the tissue are localization at the plasma membrane either non-polar or polar fashion(Péret *et al*., 2012; Swarup and Bhosale, 2019). The cellular level of auxin and the direction of auxin flow are determined by the combined activities of the PINs, ABCB, and AUX/LAX transporters (Kierzkowski *et al*., 2013).

In auxin research, the synthetic auxin transport inhibitors has been extensively used as tools to understand the role of auxin transport in cellular functions. Several auxin transport inhibitors have been characterized so far, but naphthylphthalamic acid (NPA) is the most commonly auxin efflux transport inhibitor used in the auxin researches (Teale and Palme, 2018). Although the exact mode of NPA is unclear, NPA-mediated auxin transport inhibition could associated with its direct interaction with PNA or ABCB transporters.

The capacity regeneration of plants from explants is a major importance for or biotechnological breeding such as plant propagation of elite cultivars and genetic engineering. Applying the exogenous auxin in plant tissue culture systems is a critical factor for plant regeneration. The natural auxin indole-3-acetic acid (IAA) or the other natural auxin such as 4-chloroindole-3-acetic acid (4-Cl-IAA), ndole-3-butyric acid (IBA) and phenylacetic acid (PAA) are less effective for *in vitro* plant regeneration in compered with synthetic auxins such as 2,4-dichlorophenoxyacetic acid (2,4-D) and naphthylene acetic acid (NAA). Among synthetic auxins, 2,4-D is the most effective auxin that is commonly used for plant *in vitro* regeneration. However, it remains unclear, why 2,4-D is more efficient compared to other auxin analogs. Although 2,4-D mimics the auxin activity of IAA at the molecular level (auxin signaling) (Pufky *et al*., 2003; Tan *et al*., 2007), in contrast to a rapid reduction IAA level via conjugation and degradation mechanisms in plant cells, 2,4-D is more stable (Eyer *et al*., 2016). In addition, 2,4-D is less efficient substrate for PIN and ABCB transports, which this leads a significant accumulation of 2,4-D than IAA in plant cells (Yang and Murphy, 2009). Thus, the stability or accumulation of 2,4-D can be considered as possibility why 2,4-D is more efficient for *in vitro* regeneration compared to other auxin analogs.

In this research, we found that less effective of natural auxins for regeneration is highly associated with their low accumulation in plant cells due to cell-to-cell transport. We showed that reducing auxin efflux by addition of the auxin transport inhibitor NPA allowed natural auxins and other synthetic analogs to induce SE or improve de novo shoot regeneration capacity. These our findings can be develop into effective protocols for plant regeneration.

## Result and discussion

### Enhancement of SE capacity by auxin transport inhibition

SE is a process in which in plant somatic cells are reprogrammed to embryonic cells that subsequently develop into embryos. Many SE protocols rely on the exogenous application of the synthetic auxin analog 2,4-D, whereas the IAA, 4-Cl-IAA or IBA are not or less effective in plant regeneration. In view of high intercellular accumulation of 2,4-D compared to IAA, we hypothesized that the low accumulation of IAA in plant cells might associated with its disability in inducing SE. In Arabidopsis, immature zygotic embryos (IZEs) are the most competent tissues for SE in response to the 2,4-D (Gaj, 2001). To test our hypothesis, we examined the effect of IAA accumulation on inducing embryonic callus from Arabidopsis IZEs explants by addition of auxin transport inhibitor NPA to embryonic cultures. Interestingly, we found that addition of NPA to medium allows IAA to induce SE with similar efficiencies as 2,4-D (Fig. 1A,B), whereas IZEs incubated with IAA without NPA or NPA without IAA only produced a few embryos (Fig. 1A,B). These results indicate that the less effective of IAA in inducting SE is associated with its low accumulation due to high transport dynamic in plant cells.

**Figure 1.**
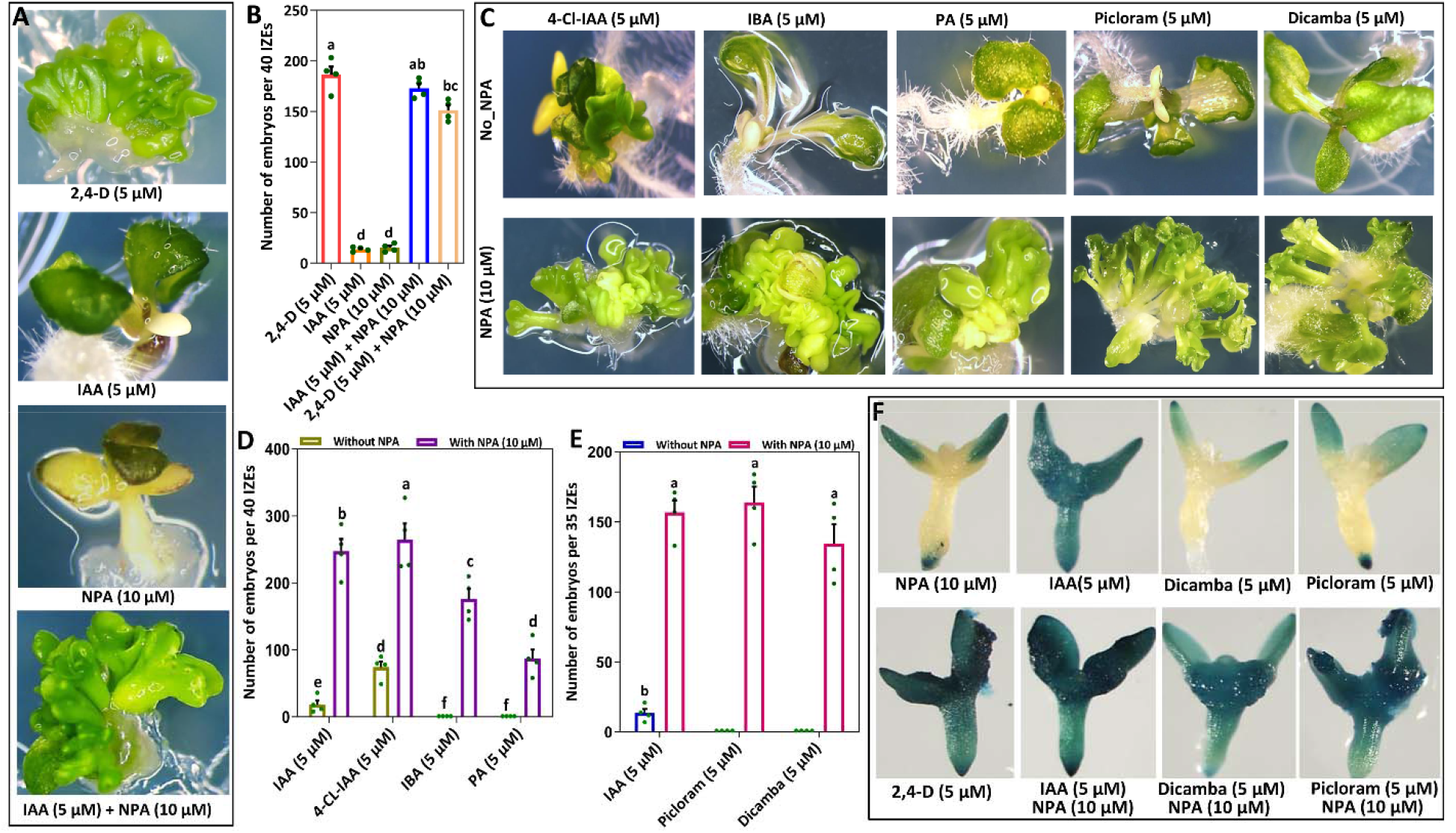
The auxin transport inhibitor NPA enhances the SE- and auxin response inducing capacity of natural and synthetic auxins in Arabidopsis. (A) The typical phenotype of an Arabidopsis immature zygotic embryo (IZE) grown for two weeks on medium supplemented with 2,4-D, IAA, NPA, or IAA and NPA, and subsequently cultured for 1 week on hormone-free medium for embryo development. (B) The number of somatic embryos per 40 IZEs that were grown for two weeks on medium supplemented with 2,4-D, IAA, NPA, IAA and NPA or 2,4-D and NPA, and subsequently cultured for 1 week on hormone-free medium. (C) The phenotype of somatic embryos formed on cotyledons of IZEs that were grown for two weeks on medium supplemented with 4-CL-IAA, IBA, PA, picloram or dicamba without NPA (upper panel) or with NPA (lower panel), and subsequently cultured for 1 week on hormone-free medium. (D, E) The number of somatic embryos per 40 IZEs (D) or 35 IZEs (E) that were grown for two weeks on medium supplemented with IAA, 4-CL-IAA, IBA, PA (D) or with IAA, picloram and dicamba (E) without NPA or with NPA, and subsequently cultured for 1 week on hormone-free medium. The dots in B, D and E indicate the number somatic embryos produced per 40 or 35 IZEs (n=4 biological replicates), bars indicate the mean and error bars indicate s.e.m. and different letters indicate statistically significant differences (P < 0.001) as determined by one-way analysis of variance with Tukey’s honest significant difference post hoc test. (F) Expression pattern of the *pDR5:GUS* reporter in Arabidopsis IZEs cultured for 7 days on medium supplemented with NPA or 2,4-D, or with IAA, picloram or dicamba without or with NPA.

To determine the impact of different level of auxin accumulation on SE, we examined the effect of different NPA concentrations (2, 5, 10, 20, 40 and 100 µM) or at present 4 µM IAA or different IAA concentrations (0.5, 1.5, 3, 4.5, 10 and 15 µM) at present of 10 µM NPA on SE. The results showed that low and high level of NPA or IAA treatments resulted in a significant reduction in the number of somatic embryos (Supplementary fig. 1A and B). To further confirm the relationship between auxin accumulation level and SE, we also analyzed the effect different concentrations of 2,4,D (0.1, 0.5, 1, 2, 5 and 10 µM) at present of 10 µM NPA on SE. The NPA treatment promotes the number of embryos at low 2,4-D concentrations (0.1, 0.5, 1, 2 µM) (Supplementary fig. 1C), whereas reduced the number of embryo at higher concentrations of 2,4,D (5, 10 µM). Together, these results indicates that the increase in the auxin accumulation promotes SE until a certain level and above of this level negatively influences SE.

To determine whether NPA treatment can promotes SE in the other natural and synthetic auxins, we analyzed the effect of NPA on capacity of SE in three other natural auxins (4-CL-IAA, IBA, PA) and two other synthetic auxins (picloram and dicomba). Intestinally, NPA treatment strongly enhanced the capacity of somatic embryo induction by 4-CL-IAA, IBA, PA (Fig. 1C,D), picloram and dicomba (Fig. 1C,E). These results indicate that the less effective of these natural and synthetic auxins in inducing of SE also contribute to their intracellular accumulation.

To examine the effect of NPA on natural or synthetic auxins-induced response at IZE explants, we employed the auxin-responsive *DR5:GUS* reporter that extensively used to visualize auxin response in Arabidopsis tissues. As expected, NPA treatment results in a relative activity *DR5:GUS* at IZE cotyledon tissues. By contrast NPA highly led to a strong *DR5:GUS* activity at present IAA, picloram, and dicomba with similar 2,4-D (Fig. 1F). These results indicate that transport inhibition of exogenesis applied natural or synthetic auxins by NPA leads their intracellular accumulation and subsequently a strong auxin response which is required for promoting SE.

### PIN- and ABCB-mediated auxin efflux reduces the capacity of IAA to induce SE

Auxin efflux is mainly facilitated by polar localization of PIN proteins on the plasma membrane. To show whether PIN carries are responsible for low efficiency of IAA-induced SE, first we examined the expression of *PIN1:PIN1-GFP, PIN2:PIN2-VENUS, PIN3:PIN3-GFP, PIN4:PIN4-GFP*, and *PIN7:PIN7-GFP* reporters in IZEs treated with IAA and IAA/NPA. Of these reporters, only PIN1-GFP and PIN2-VENUS expression was detected in IZE cotyledon tissues (Fig. 2A, B). In two-and five-day-old IZE explants, PIN1-GFP signals was not detected in IZE cotyledon tissues (Fig. 2A),while PIN2-VENUS signals was detected at the abaxial side of the cotyledons IZEs (Fig. 2B). The earliest PIN1-GFP signals was detected at the abaxial side of the cotyledons at 7 days of cultures (Fig. 2A), while PIN2-VENUS significantly decreased in cotyledons (Fig. 2B). These results indicate PIN1 and PIN2 are likely responsible IAA depletion in the cotyledon epidermal cells. We did not observed the polar localization of PIN1-GFP and PIN2-VENUS signals in the cotyledon epidermal cells (Fig. 2C), therefore we suggest that the auxin distribution in cotyledon cells by PIN1and PIN2 is likely processed in a polar auxin transport-independent manner. To investigate effect of the PIN carriers on the capacity of SE, we assessed the effect of *pin2* mutant on the capacity embryo induction by IAA and IAA/NPA. Our experiments showed that *pin2* IZEs produced significantly more number of embryo than wild type IAA/NPA treatments, whereas it has no effect the capacity of SE by IAA at absent of NPA (Fig. 2D). To know whether increase in the number embryos in *pin2* mutant is related to the depletion of IAA by PIN2 in the cotyledon cells on day 1-4 of culture, the *pin2* and wild type IZEs first were incubated on medium containing IAA without NPA for 4 days then were transformed to medium containing IAA with NPA. Although the early incubation of IZEs to medium containing only IAA led to significant decrease the number embryos of the *pin2* and wild type IZEs (Fig. 2E), *pin2* IZEs was less sensitive to this early incubation than wild type (Fig. 2E). Therefore increase in the number embryos in *pin2* mutant is associated to less depletion of IAA in the cotyledon epidermal cells on 1-4 day-old IZEs.

**Figure 2.**
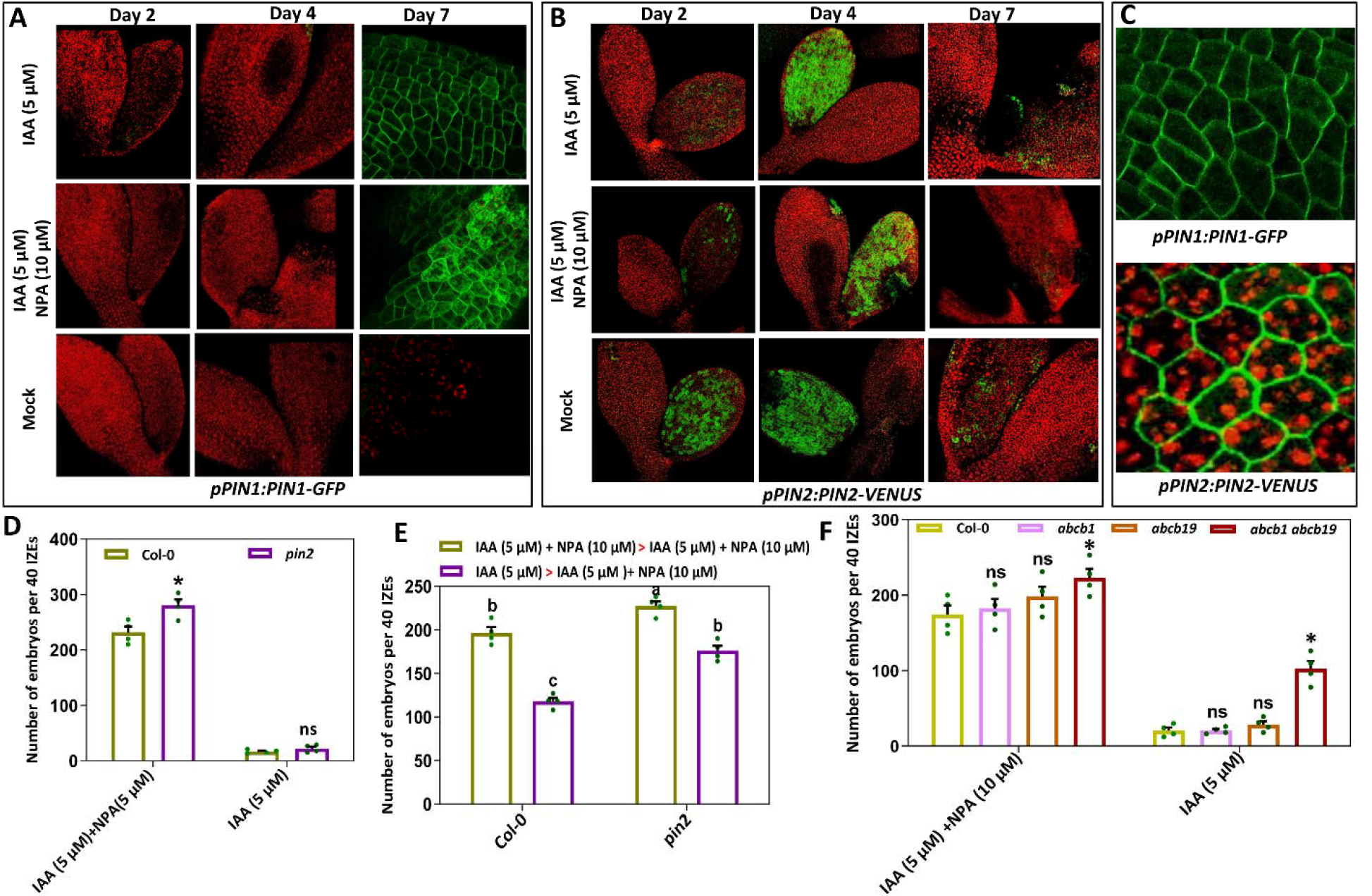
PIN- and ABCB-mediated auxin efflux reduces the capacity of IAA to induce SE. (A, B) The expression patterns of *pPIN1:PIN1-GFP* (A) and *pIN2:PIN2-VENUS* (B) in cotyledon of IZEs cultured for two, four or seven days on medium supplemented with IAA (upper panel), with IAA and NPA (middle panel), or without any addition (mock, lower panel). (C) Non-polar localization of PIN1-GFP (upper panel) and PIN2-VENUS (lower panel) in cotyledon epidermis cells of IZEs after respectively seven and two days of culture. (D) The number of somatic embryos per 40 wild-type or or *pin2* mutant IZEs that were first grown for two weeks on medium supplemented with IAA or IAA and NPA, and subsequently cultured for 1 week on hormone-free medium for embryo development. (E) Differential reduction in the number of somatic embryos formed on *pin2* IZEs compared with wild type which were first grown for four days on medium supplemented with IAA and then transferred in to medium supplemented IAA/NPA for 10 days. (F) The number of somatic embryos per 40 IZEs that were first grown for two weeks on medium supplemented with IAA, and IAA/NPA, and subsequently cultured for 1 week on medium without IAA and NPA at wild type, *abcb1, abcb19, abcb1 abcb19* mutants. In D-F the dots indicate the number somatic embryos produced per 40 IZEs (n=4 biological replicates), bars indicate the mean and error bars indicate s.e.m. and the asterisk indicates a significant difference with wild-type (P<□0.01) as determined by a two-sided Student’s *t*-test.

To find out whether the other auxin efflux transports might also be involved in reduced capacity of IAA-induced SE, we assessed the capacity of somatic embryo induction by IAA and IAA/NPA in *ABCB1, ABCB19* and *ABCB1 ABCB19* mutants. The single mutant *ABCB1, ABCB19* did not showed significantly different in the number of embryo induced by IAA/NPA and IAA than wild (Fig. 2F), whereas double mutant *ABCB1 ABCB19* IZEs promotes SE in both IAA/NPA and IAA treatments (Fig. 2F). This result indicates that ABCB-mediated auxin efflux also reduces the capacity of IAA to induce SE.

### Natural auxin-induced somatic embryos show improved seedling conversion

At stage of convention somatic embryo to seedling, we noticed efficient shoot development from embryos induced by the natural auxins in the presence of NPA than 2,4-D (Supplementary fig. 2). To monitor this remarkable embryo convention, we first isolated the single full-developed embryos (Fig. 3A) induced by the natural auxins or 2,4-D, then we assessed the convention of these single embryos transferred into new medium. Importantly, all somatic embryos induced by the natural auxins in the presence of NPA showed a normal embryo to seedling conversion (Fig. 3C,D), whereas for the 2,4-D system this was limited to 50-60% of the embryos (Fig. 3B,D) with less synchronized growth of shoot among seedlings (Fig. 3B) and less synchronized flowering timing of plants in soli (Supplementary fig. 3).

**Figure 3.**
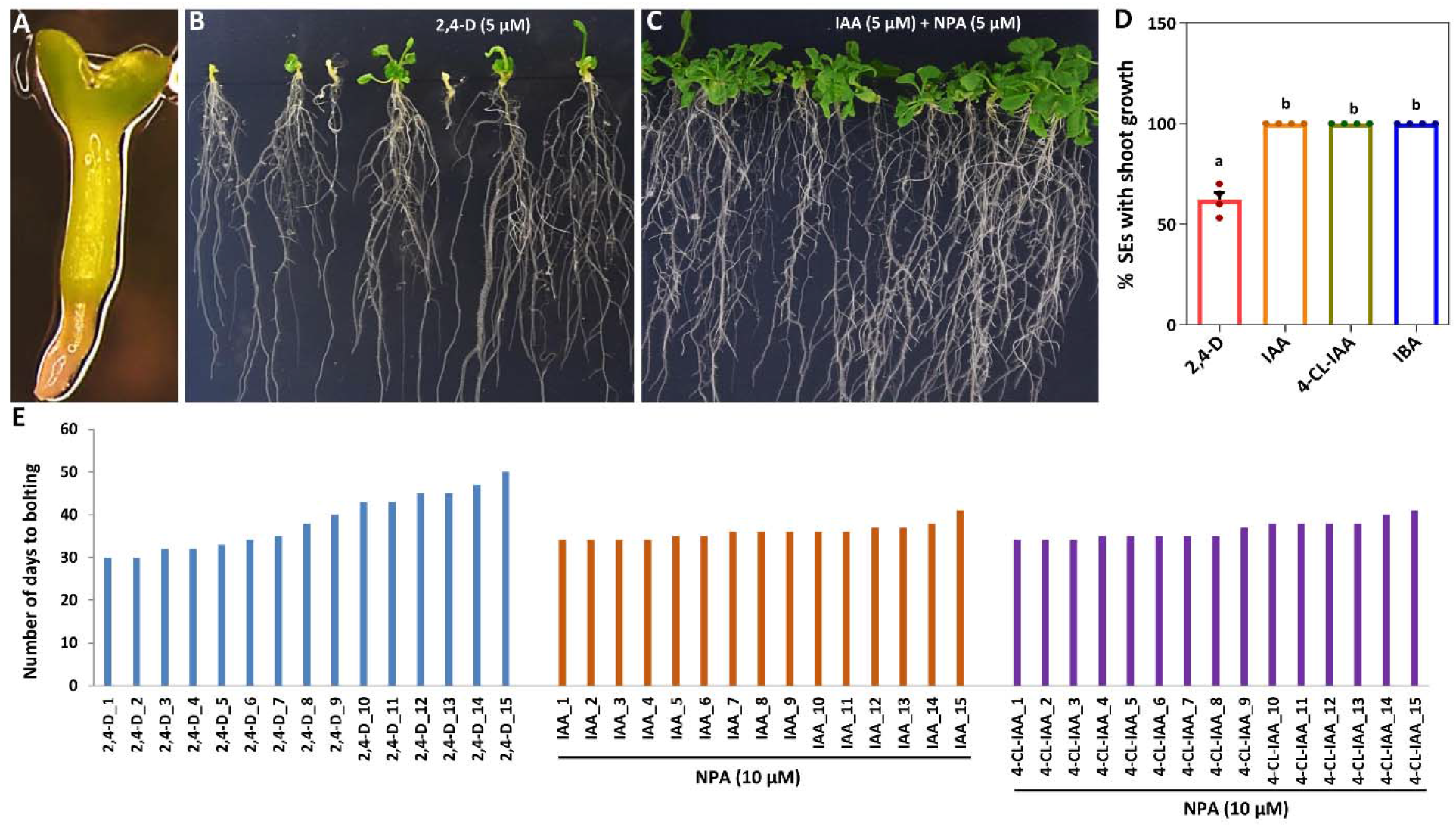
Natural auxin-induced somatic embryos show improved seedling conversion. (A) The phenotype of a single natural auxin or 2,4-D induced somatic embryo cultured on hormone-free medium for conversion to seedling. (B,C) The phenotype of seedlings derived from somatic embryos induced on 2,4-D (B) or IAA and NPA containing medium (C) after germination for 2 weeks on hormone-free medium. (D) The embryo to seedling conversion rate of somatic embryos induced by 2,4-D, IAA and NPA, 4-CL-IAA and NPA or IBA and NPA 2 weeks after germination on hormone-free medium. The dots indicate the percentage of 20 somatic embryos showing proper embryo to seedling conversion (n=4 biological replicates), bars indicate the mean and error bars indicate s.e.m. and different letters indicate statistically significant differences (P < 0.01) as determined by one-way analysis of variance with Tukey’s honest significant difference post hoc test.

We also assessed the convention efficiency of somatic embryos induced from *Camelina sativa* IZEs by 2,4-D in compared with 4-Cl-IAA/NPA. Similar to Arabidopsis, the most of somatic embryos induced by 4-Cl-IAA showed a normal embryo to seedling conversion (Fig. 3F,G), whereas for the 2,4-D was limited to 30-40% of the embryos (Fig. 2E,G).

Low rate of somatic embryo convention induced by 2,4-D and high variation in the growth pattern among plants derived-somatic embryos, has been reported in many plant species (Garcia *et al*., 2019). Based on these data we conclude, that the efficiency of SE-based plant propagation can be significantly improved by applying the natural auxins in the presence of the auxin transport inhibitor NPA.

### Improvement of de novo shoot regeneration capacity by auxin transport inhibition

De novo shoot regeneration is a two-step regeneration process: inducing pluripotent callus from explants on a auxin-rich callus-inducing medium (CIM) then inducing apical meristems from the callus on a cytokinin-rich shoot-inducing medium (SIM). To determine whether de novo shoot regeneration can also be improved by applying natural or synthetic auxins in the presence of NPA, the capacity of shoot regeneration were tested in callus induced in hypocotyl and cotyledons of *Brassica napus* seedlings by 2,4-D and 4-CL-IAA in the presence of NPA. The analysis showed that present of NPA in 2,4-D-CIM significantly decreases and increases the capacity of shoot regeneration of callus from hypocotyls and cotyledons respectively (Fig. 4A-C), while addition of NPA to 4-CL-IAA-CIM resulted in very effective regeneration of callus from both hypocotyls and cotyledons (Fig. 4A-C). These results indicate that the auxin transport inhibition can also promote the regeneration capacity of callus induced either by synthetic or natural auxins.

**Figure 4.**
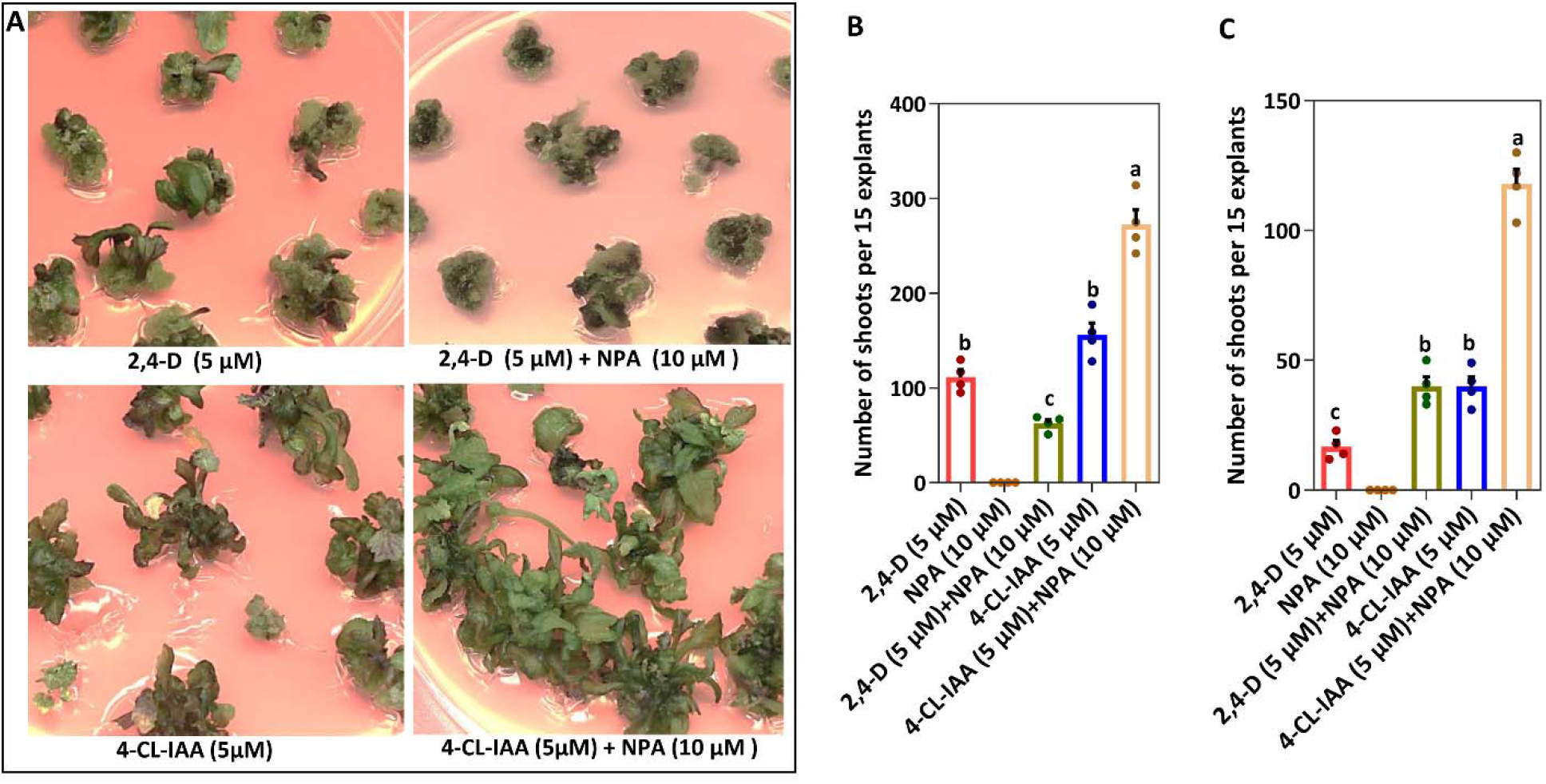
The auxin transport inhibitor NPA enhances de novo shoot regeneration capacity in *Brassica napus*. (A) The phenotype of shoot formed on callus induced hypocotyls for 10 days on medium supplemented with 2,4-D, 2,4-D/NPA, 4-Cl-IAA or 4-Cl-IAA /NPA, and subsequently cultured for 3 weeks on medium supplemented with 10 µM 6-Benzylaminopurine (BA) for shoot regeneration. (B,C) The number of shoots formed on callus induced from 15 hypocotyl (B) or 15 cotyledons (C) explants for 10 days on medium supplemented with 2,4-D, NPA, 2,4-D with NPA, 4-Cl-IAA or 4-Cl-IAA with NPA, and subsequently cultured for 3 weeks on medium supplemented with 10 µM BA for shoot regeneration. The dots in B and C indicate the number of shoots produced per 15 explants (n=4 biological replicates), bars indicate the mean and error bars indicate s.e.m. and different letters indicate statistically significant differences (P < 0.001) as determined by one-way analysis of variance with Tukey’s honest significant difference post hoc test.

We also noticed early appearing of shoot buds on callus induced by 4-CL-IAA than 2,4-D (Supplementary fig. 4) which this can allows to development a faster regeneration protocol for *B. napus* by natural auxins. In addition, we observed a better elongation of shoots formed on callus induced by 4-CL-IAA than 2,4-D. To monitor this, we isolated the shoot buds formed on callus induced by 4-CL-IAA than 2,4-D, then we assessed the growth of these shoots in a free hormone medium. Importantly, the shoots induced by 4-CL-IAA showed synchronized growth than 2,4-D (Supplementary fig. 5). Based on these data we conclude, that the efficiency of shoot regeneration-based plant propagation can be significantly improved by applying the natural auxins.

## Materials and methods

### Plant material and growth conditions

All *Arabidopsis thaliana* lines used in this study were in the Columbia (Col-o) background. The transgenic lines *pDR5:GFP* (Ottenschläger *et al*., 2003), *pWOX2:NLS-YFP* (Breuninger *et al*., 2008), *pPIN1:PIN1-YFP* (Benkova *et al*., 2003) and *pPIN2:PIN2-VENUS* (Blakeslee *et al*., 2007) have been described previously. *abcb1, abcb19*, and *abcb1abcb19* plant lines were obtained from the Nottingham Arabidopsis Stock Centre (NASC). Seeds were sterilized in 10 % (v/v) sodium hypochlorite for 12 minutes and then washed four times in sterile water. Sterilized seeds were plated on half MS medium (Murashige and Skoog, 1962) containing 1 % (w/v) sucrose and 0.7 % agar. Seedlings, plants, and explants were grown at 21°C, 70% relative humidity and 16 hours photoperiod.

### Somatic embryogenesis

For the isolation of IZEs at the bent cotyledon stage of development, siliques were harvested 10-12 days after pollination, sterilized in 10 % (v/v) sodium hypochlorite for 7 minutes and then washed four times in sterile water. IZEs were dissected from the siliques inside a laminar flow cabinet (Gaj, 2001). For induced SE, IZEs were cultured on solid B5 medium supplemented with 2,4-D, IAA, 4-Cl-IAA, IBA, PA, picloram and dicomba with or without NPA, 2 % (w/v) sucrose and 0.7 % agar (Sigma) for 2 weeks. Subsequently, the embryonic structures were allowed to develop further by transferring the explants to half MS medium with 1 % (w/v) sucrose and 0.7 % agar (Sigma) without hormones. One week after subculture, the capacity to induce SE was scored under a stereomicroscope as the number of somatic embryos produced from IZEs per plate. Four plates were scored for each experiment.

### Shoot regeneration

Brassica napus cultivar wstar was used in this study. Seeds were sterilized in 10 % (v/v) sodium hypochlorite for 12 minutes and then washed four times in sterile water. Sterilized seeds were plated on half MS medium containing 1 % (w/v) sucrose and 0.7 % agar without plant growth regulators. Hypocotyl explants were excised from 7-day-old seedlings and cultured on MS solid media supplemented with 1% sucrose and with 2,4-D, 4-Cl-IAA with or without NPA. The explants were transferred into MS solid media supplemented 6-Benzylaminopurine (BA) for shoot regeneration. 3 weeks after transferring explants, the capacity shoot regeneration was scored under a stereomicroscope as the number of number shoots produced from hypocotyl per plates. Four plates were scored for each experiment.

### GUS Staining

Histochemical staining of transgenic lines expressing the β-glucuronidase (GUS) reporter for GUS activity was performed as described previously (Anandalakshmi et al., 1998) for 4 hours at 37 °C, followed by rehydration in a graded ethanol series (75, 50, and 25 %) for 10 minutes each.

### Microscopy

GUS-stained tissues and cultured IZE explants were observed and photographed using a LEICA MZ12 microscope (Switzerland) equipped with a LEICA DC500 camera.

Confocal Laser Scanning Microscopy (CSLM) was performed with a ZEISS-003-18533. GFP and YFP were detected using a 534 nm laser, a 488 nm LP excitation filter and a 500-525 nm band pass emission filter.

Simultaneously, background fluorescence (of e.g. chlorophyll) was detected with a 650nm long pass emission filter. Images were captured with ZEISS ZEN2009 software. Unmodified images were cropped (if needed) and used for assembly into figures in MS Powerpoint. Assembled figures were saved as pdf files and converted to tif files in Adobe Photoshop.

---

**Supplementary figure 1.**
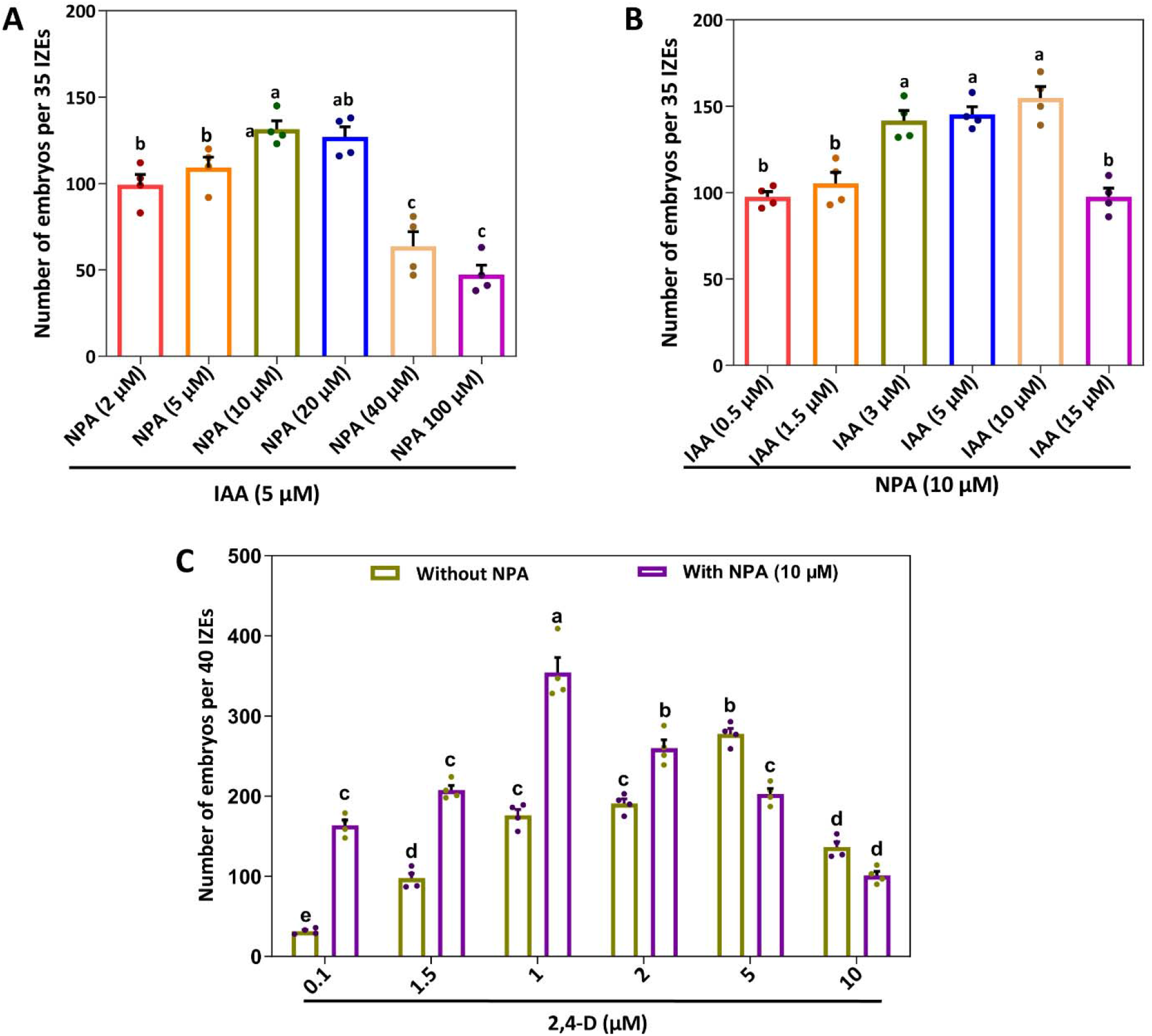
Inhibition of auxin transport lowers the exogenous auxin concentration required for efficient SE induction. (A) Effect of different concentrations of NPA on the capacity to induce somatic embryos on IZEs cultured on medium with IAA. (B) Effect of different concentrations of IAA on the capacity to induce somatic embryos on IZEs cultured on medium with NPA. The dots in A, B indicate the number somatic embryos produced per 35 IZEs (n=4 biological replicates), bars indicate the mean and error bars indicate s.e.m. and different letters indicate statistically significant differences (P < 0.01) as determined by one-way analysis of variance with Tukey’s honest significant difference post hoc test. (C) Effect of different concentrations of 2,4-D on the capacity to induce somatic embryos on IZEs cultured on medium with and without NPA. The dots indicate the number somatic embryos produced per 40 (n=4 biological replicates), bars indicate the mean and error bars indicate s.e.m. and the asterisk indicates a significant difference (P<□0.01) as determined by the two-sided Student’s *t*-test.

**Supplementary figure 2.**
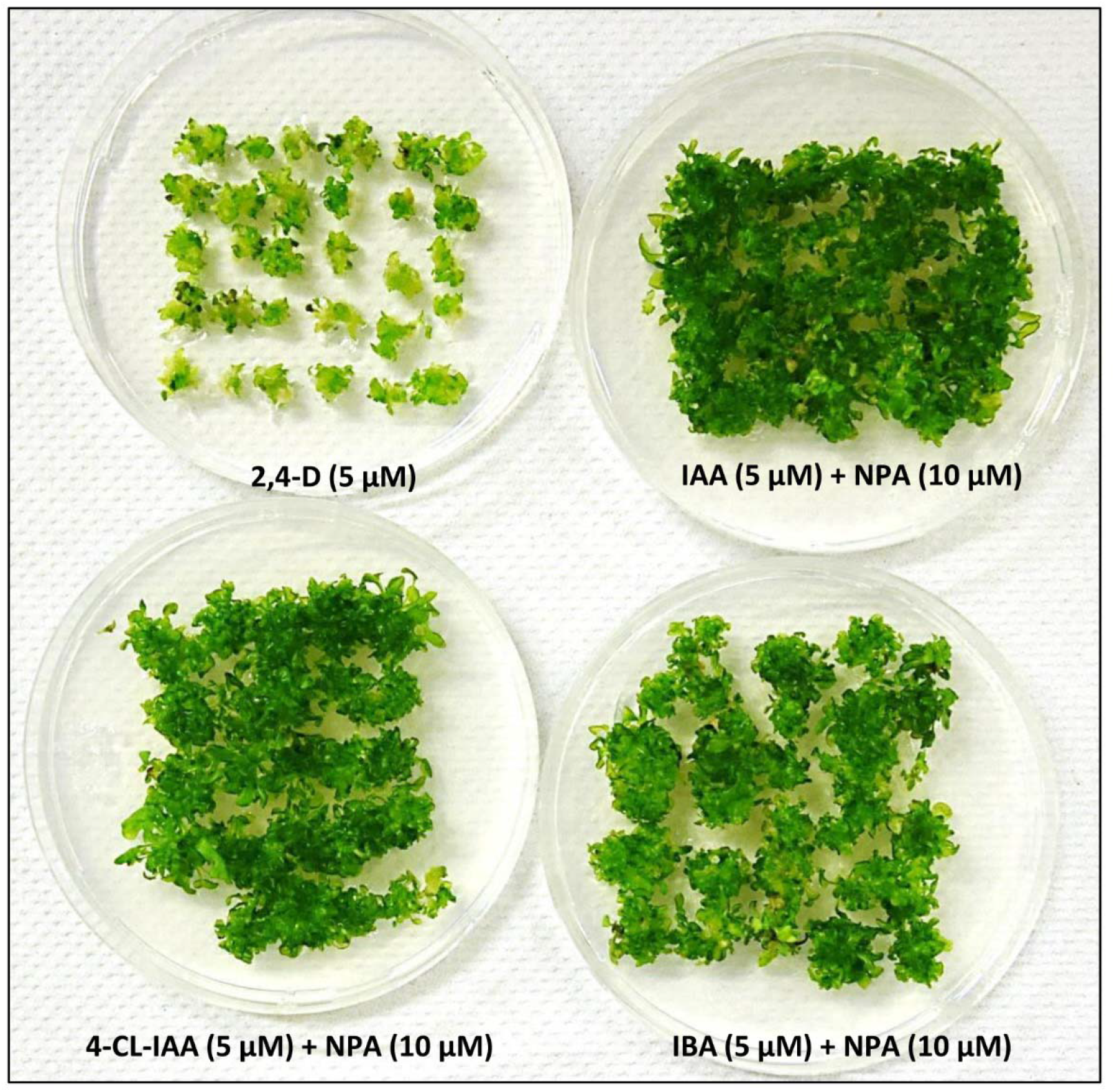
Natural auxin-induced somatic embryos show improved seedling conversion. The phenotype of shoot derived from somatic embryos induced on 2,4-D, IAA/NPA, 4-CL-IAA/NPA or IBA/NPA containing medium after conversion for 2 weeks on hormone-free medium.

**Supplementary figure 3.**
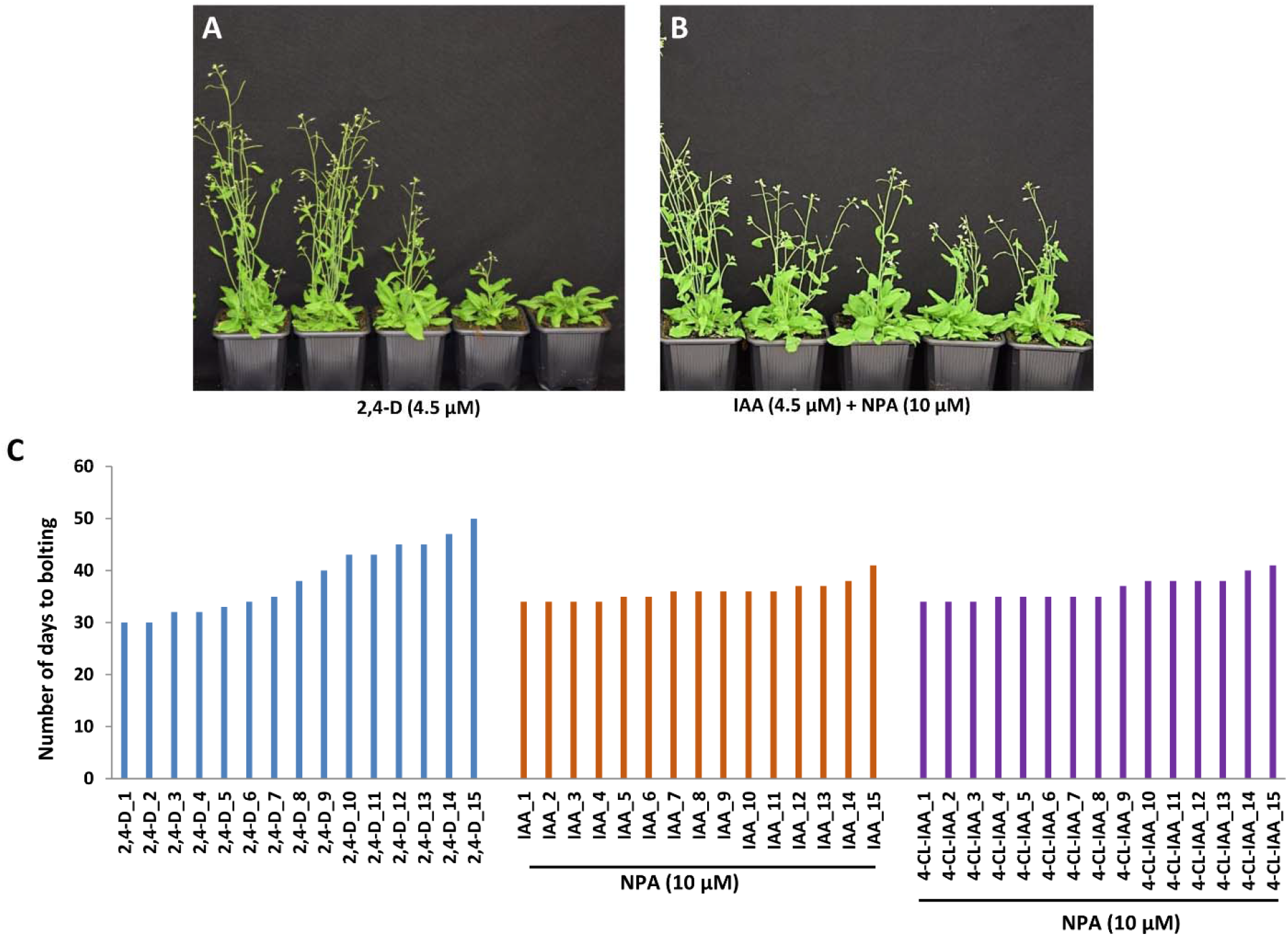
Natural auxin-induced somatic embryos show improved plant development. (A, B) The shoot phenotypes of 45-day-old plants derived from somatic embryos induced by 2,4-D (A) and IAA/NPA (B). (C) The number of days to bolting in plants derived from somatic embryos induced by 2,4-D, IAA/NPA, and 4-CL-IAA/NPA.

**Supplementary figure 4.**
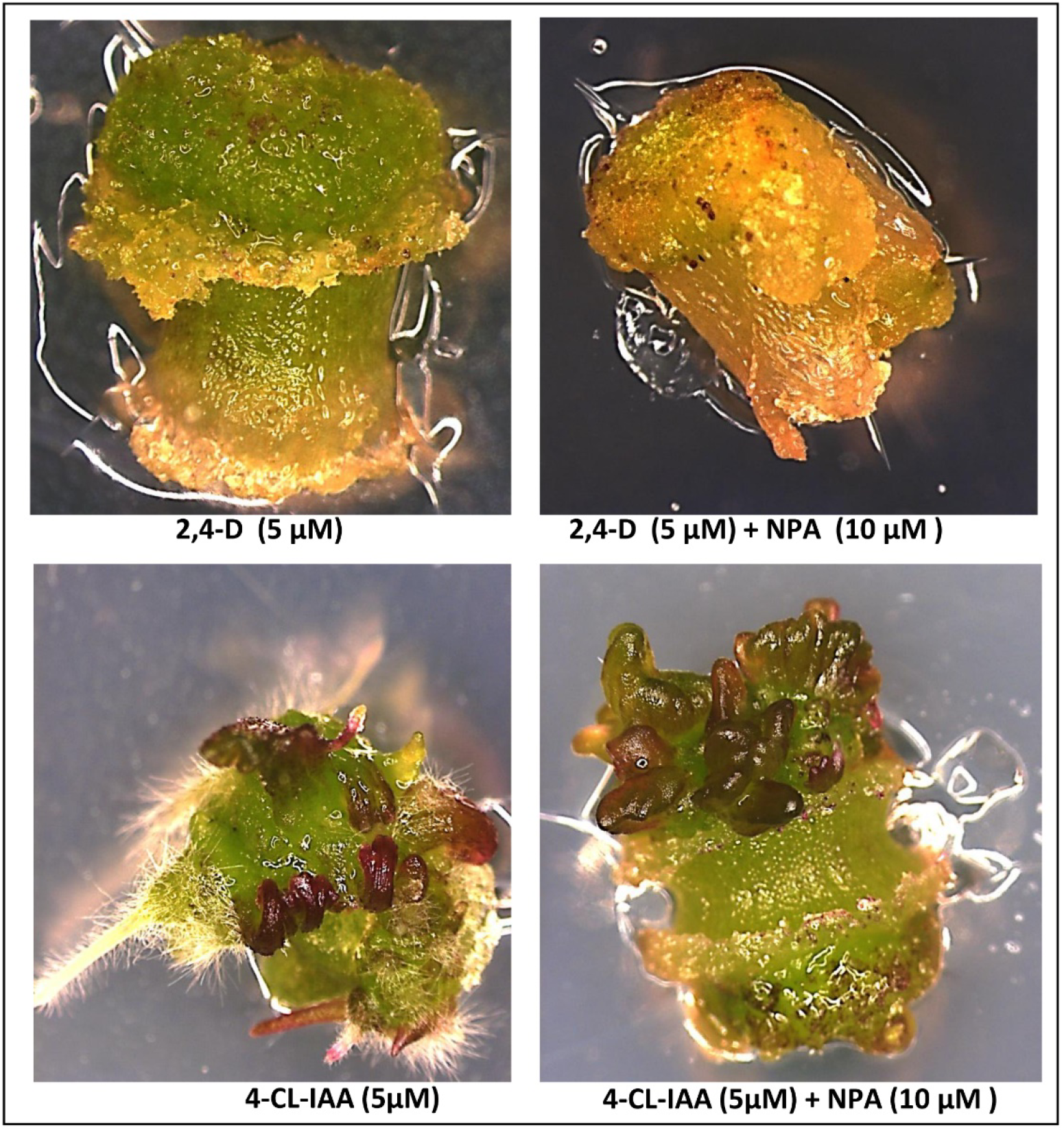
Acceleration in shoot buds formation on callus induced by 4-CL-IAA. The phenotype of callus induced from hypocotyls by 2,4-D, 2,4-D/NPA, 4-Cl-IAA or 4-Cl-IAA /NPA, and subsequently cultured for 1 week on medium supplemented with 10 µM 6-Benzylaminopurine (BA).

**Supplementary figure 5.**
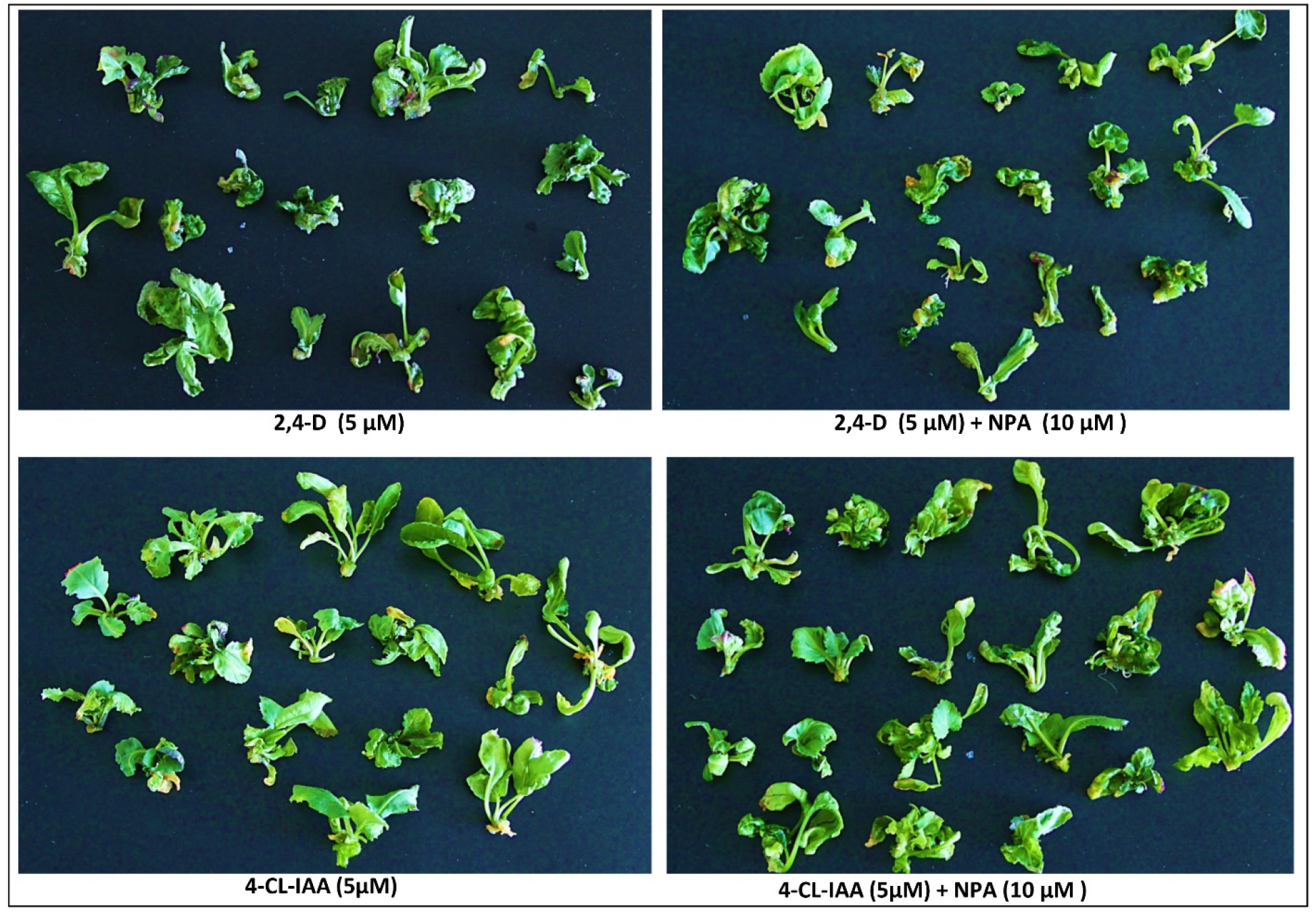
The shoot buds formed on callus induced by 4-CL-IAA shows synchronized growth than 2,4-D. The phenotype of shoot derived from shoot buds formed on callus by 2,4-D, 2,4-D/NPA, 4-Cl-IAA or 4-Cl-IAA /NPA, for 2 weeks on hormone-free medium.

